# Correcting for population stratification reduces false positive and false negative results in joint analyses of host and pathogen genomes

**DOI:** 10.1101/232900

**Authors:** Olivier Naret, Nimisha Chaturvedi, Istvan Bartha, Christian Hammer, Jacques Fellay

## Abstract

Studies of host genetic determinants of pathogen sequence variation can identify sites of genomic conflicts, by highlighting variants that are implicated in immune response on the host side and adaptive escape on the pathogen side. However, systematic genetic differences in host and pathogen populations can lead to inflated type I (false positive) and type II (false negative) error rates in genome-wide association analyses. Here, we demonstrate through simulation that correcting for both host and pathogen stratification reduces spurious signals and increases power to detect real associations in a variety of tested scenarios. We confirm the validity of the simulations by showing comparable results in an analysis of paired human and HIV genomes.

## 1 Introduction

Important inter-individual differences can be observed in human responses to infections, and in recent years researchers have started to explore the genetic underpinning of this phenotypic diversity [1,2,3,4,5]. A better understanding of host-pathogen interactions at the genomic level could help explain pathogenesis, predict disease outcome or develop new therapeutics or vaccines.

Multiple genome-wide association studies (GWAS) of clinical outcomes have identified human genetic variants that play a modulating role in infectious diseases [6,7,8,9]. One of the most prominent examples is the strong association between human leukocyte antigen (HLA) variation and HIV-1 control [10,11,12]. To further explore the potential impact of human genetic diversity on infection, we recently proposed to integrate host and pathogen genomic data in a single analytic framework (which we called genome-to-genome analysis, or G2G) [13]. Through a pilot study in an HIV-infected population, we demonstrated that it is possible to detect the sites of genomic conflict between the host and the pathogen. Host restriction factors can thus be uncovered by identifying the escape mutations that accumulate in the pathogen genome in response to selection pressure exerted by host genetic variants.

Systematic genetic differences in a study population, or population stratification, can lead to inflated type I (false positive) and type II (false negative) error rates [14,15,16]. In standard GWAS, the association analyses are corrected for stratification by adding host covariates, such as principal components calculated from the host genomic data, to the model [17,18,19,20,21]. However, accounting for population structure becomes more challenging in a G2G analysis, when these systematic differences can be present in both the host and the pathogen populations.

In this paper, we explore the effects of population stratification in G2G analyses. We first present a basic introduction to the G2G framework and to the methods used for population stratification correction. We then simulate host and pathogen variation using a broad array of parameters including stratification on both sides, which allows us to pinpoint the various effects of population stratification on different statistical models. We finally test for associations between genome-wide human genotypes and HIV-1 sequence diversity in a real-life dataset. The simulation models, as well as all the steps for analyzing the simulated datasets are implemented in R, and the framework is available on GitHub [22].

## 2 Material and Methods

### 2.1 Genome-to-genome framework

In our G2G analysis, we use a regression model to search for associations between host and pathogen genetic variation. Host genetic variation is represented by a genotype data matrix with n samples and p single nucleotide polymorphisms (SNPs). Pathogen variation is represented by a matrix with n samples and m binary variants. Then, if the variation data corresponding to a single position of the pathogen genome *y* with n observations is *y* = (*y*_1_,*y*_2_,…,*y*_*n*_)^T^, the regression model can be written as:

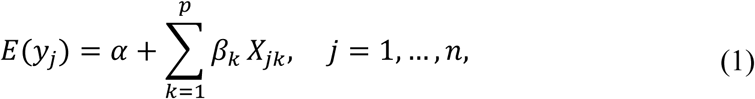

where *X*_*jk*_ is the genotype for the*j*^*th*^ sample and the *k*^*th*^ SNP, and *α* is the intercept. *β*_*κ*_ is the coefficient value for the *k*_*th*_ SNP; it explains the association patterns between the pathogen variations ***y*** and genotype data {*x*_*jk*_, *j* = 1,…,*n*; *k* = 1,…,*p*}. Once all the *β* coefficients are known, the association p-values are obtained by testing the null hypothesis *H*_0_:*β* = 0 against the alternative hypothesis *H*_1_*β* ≠ 0.

### 2.2 Simulation study

#### 2.2.1 Generation of host genetic data (SNPs)

Our simulation design is based on the Balding Nicholas Model [23], which provides a framework for estimating the probabilities of observed genotypes, taking into account population structure and variance in allele frequency estimates. For simulating stratification between host subpopulations, we start with drawing an ancestral allele frequency *R*_*ƒ*_ for each SNP from the uniform distribution on [0.1,0.9]. To generate a SNP stratified between two subpopulations we use a specified Wright’s F-statistics (*F*_*ST*_) [14, 24] for drawing two alternate allele frequencies 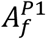 and 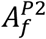from a *β* distribution with parameters:

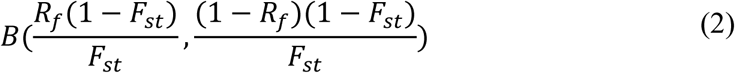

This distribution has mean *R*_*ƒ*_ and variance *F*_*ST*_*R*_*ƒ*_(1 -*R*_*ƒ*_). Then, for a population P1 with an alternate allele frequency of 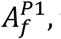 the genotypes 0 (homozygous for reference allele), 1 (heterozygous) or 2 (homozygous for alternate allele) are assigned to each sample with probabilities 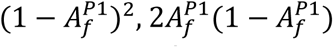 and 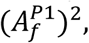 respectively. For an unstratified SNP, the genotypes 0, 1 and 2 are assigned with probabilities (1 - *R*_*ƒ*_)^2^, 2*R*_*ƒ*_(1 -*R*_*ƒ*_), and 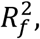respectively.

#### 2.2.2 Generation of pathogen variables

To create a binary variable of value 0 (absence) or 1 (presence) for each pathogen variant, we start by generating the background random variations represented by a binary vector, *Y*_*bg*_. We draw the parameter *R*_*ƒ*_ from a uniform distribution, which defines the ancestral mutation rate for a given position on the pathogen genome. For variables that are not stratified between groups A and B, *Y*_*bg*_ is generated from a binomial distribution with *R*_*ƒ*_ being the probability of mutation. For variables that are stratified between A and B, we use a specified *F*_*ST*_ to obtain two alternate mutation frequencies 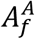 and 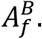 These alternate frequencies are then used as probabilities for generating *Y*_*bg*_,in groups A or B, from a binomial distribution.

In a second step, we add associations between a selected set of binary pathogen variable and certain host SNPs. We start by generating a binary vector, *Y*_*ca*_, using the logistic function given as:

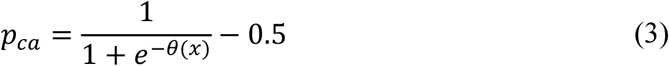

Here, *θ*(*x*) = *xβ*, with *x* representing a vector of genotypes for all samples and *β* representing the association coefficient. The vector of probabilities pca is then used to generate *Y*_*ca*_ from a binomial distribution. Using *Y*_*bg*_ and *Y*_*ca*_, the final pathogen variations are generated as:

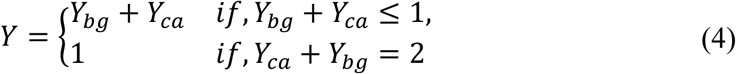

Therefore, a sample can have a value of 1 for a given pathogen variant in the presence of background variation (*Y*_*bg*_ = 1), causal variation (*Y*_*ca*_ = 1), or both (*Y*_*ca*_ = *Y*_*bg*_ = 1). This strategy allows for the generation of a matrix of binary pathogen variants that includes background variation as well as variants associated with host SNPs.

### 2.3 Joint analysis of human and HIV genetic variation

#### 2.3.1 Study participants

Human and viral genomic variation data from 1668 participants in the Swiss HIV Cohort Study [25](www.shcs.ch) were included in the analysis.

#### 2.3.2 Human genotype data

Human genome-wide genotyping data were generated in the context of previous studies. SNPs were excluded on the basis of per-individual missingness (> 3), genotype missingness (> 1), and marked deviation from Hardy-Weinberg equilibrium (*p* < 1× 10-^7^). Genotype imputation was performed on the Sanger imputation server, using EAGLE2 [26] for pre-phasing and PBWT [27] with the 1000 Genomes Phase 3 reference panel (1000 Genomes Project Consortium et al. 2015). Imputed variants were filtered based on imputation INFO score (dropped if < 0.8).

#### 2.3.3 HIV sequence data

Partial retroviral sequence data were obtained as part of routine clinical testing for resistance against antiretroviral drugs. The complete protease region (PR) and 50% of the reverse transcriptase (RT) region were sequenced from pretreatment plasma samples. We defined an amino acid residue as variable if at least 10 study samples carried an alternative allele. Per position, separate binary variables were generated for each alternate amino acid, indicating the presence or absence of that allele in a given sample.

### 2.4 Association analyses

We used logistic regression to test for associations between host SNPs and HIV amino acid variants. All models were run with plink [28], assuming an additive genetic model. To assess the effect of stratification correction, we used four different approaches. First, we tested for associations without any correction for stratification. Second, we adjusted the model for covariates capturing host stratification. Third, we adjusted the model for covariates capturing pathogen stratification. Fourth, we used both host and pathogen covariates in the model. This resulted in four distinct sets of association results. All p-values were corrected for multiple testing based on Bonferroni thresholds.

## 3 Results

### 3.1 Simulation analysis

Our simulation study includes paired host and pathogen data from 5000 samples. The hosts are stratified between two subpopulations P1 and P2 of 2500 samples each and the pathogens are stratified between two groups, A and B of 2500 samples each. To generate spurious signal, we created unequal groups of paired host and pathogen. Within P1, 1500 samples are infected by strain A and 1000 by strain B. Within P2, 1000 samples are infected by strain A and 1500 are infected by strain B. Genomic variation data include 50,300 host SNPs and 400 pathogen variants. The parameters used to generate host SNPs and pathogen variants are presented in Table 1 and 2, respectively. The first column of the tables (‘Quantity’) gives the number of host SNPs or pathogen variant. The second column (‘Causal association’) specifies the presence or absence of association between pathogen variants and host SNPs. For host SNPs, it also specifies the strength of the causal relationship (value of the β coefficient), as well as the corresponding associated pathogen variant (line reference for Table 2). The columns 3 (‘Major stratification’) and 4 (‘Minor stratification’) represent the two possible levels of stratification and specify the strength of stratification through FST values. These two columns also show which subpopulation has the highest/lowest variation frequency.

**Table 1.**
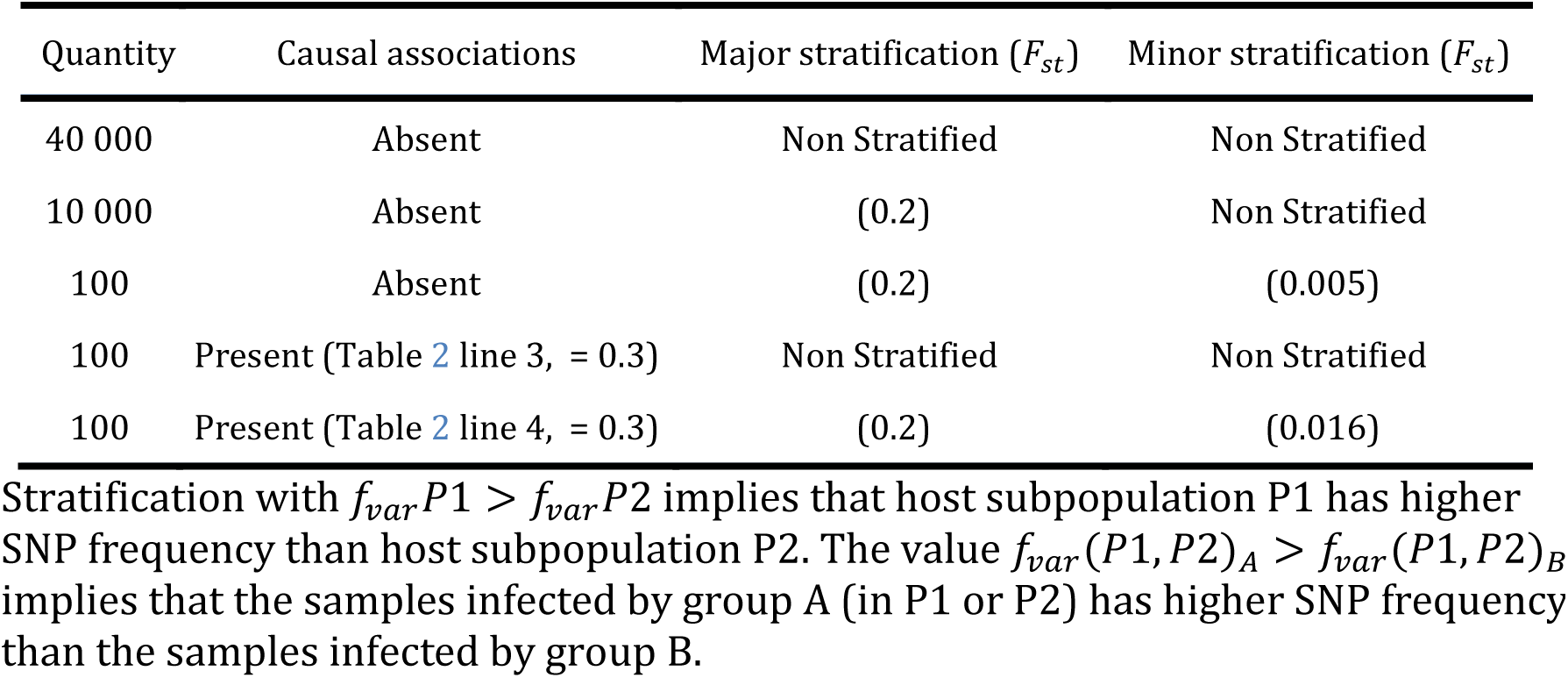
Host SNPs parameters

**Table 2.** Pathogen variants parameters

| Quantity | Causal associations | Major stratification ($F_{st}$) | Minor stratification ($F_{st}$) |
| --- | --- | --- | --- |
| 100 | Absent | (0.2) | Non Stratified |
| 100 | Absent | (0.2) | (0.005) |
| 100 | Present | Non Stratified | Non Stratified |
| 100 | Present | (0.2) | (0.01) |

Stratification defined by $f_{var}A > f_{var}B$ implies that pathogen strain A has higher pathogen variation frequency than pathogen strain B. The value $f_{var}(A, B)_{P1} > f_{var}(A, B)_{P2}$ implies that the pathogen genomes (A and B) from host population P1 exhibit a higher mutation rate than the pathogen genomes from P2.

We used logistic regression to test for association between each host SNP and pathogen variant (20,120,000 tests in total). To correct for host stratification, we used the top 5 principal components of the human genotyping data, and to correct for pathogen stratification, we used the group marker as covariates. The significance was assessed using the Bonferroni threshold of 2.49 × 10-^9^. The results are presented in the next three paragraphs: [1] ‘No spurious signals’; [2] ‘False positives’, where the only observed signals are due to stratification; [3] ‘Power gain’, where real association signals are weakened by stratification. The input parameters to reproduce similar datasets using the G2G-Simulator [22] as well as the exact simulated dataset detailed in the following section [29] are available online

#### 3.1.1 No spurious signal

A total of 17,060,100 association tests fall into this category. For host-pathogen variant pairs with no association (N=17,060,000), the median p-value is 0.5 regardless of the correction applied. For the 100 pairs of host SNPs (line 4, Table 1) and pathogen variants (line 3, Table 2) that are associated, we observe similar p-values irrespectively of the type of stratification correction, with median p-values ranging between 5.3 × 10-^7^ and 6.7 × 10-^7^ for the four different approaches of correction. So, in the absence of spurious signals, correcting for any population structure has negligible effect on the association results. The Manhattan plots of the corresponding tests are available in the <u>online supplementary materials</u> [30] by selecting ‘No spurious signal’.

#### 3.1.2 False positives

Here, we study the effects of stratification correction on false positive signals, through two scenarios, with different levels of stratification complexity.

**Scenario 1, optimal correction:** This scenario presents association tests between 10,000 host SNPs (l.2 Table 1) and 100 pathogen variants (l.1 Table 2) resulting in 1,000,000 tests (figure 1A). The top half of the figure represents host SNPs frequencies in the two populations. Host SNPs are stratified between P1 and P2 with higher SNP frequency in P1, represented by more ‘+’ signs under P1 than in P2. The bottom half of the figure represents pathogen variants frequencies in the two strains. Pathogen variants are stratified between A and B with higher variant frequency in A than B (figure 1B, more ‘+’ signs under A than B). Since there are more samples in P1 infected by pathogen group A and more samples in P2 infected by pathogen group B, it creates a correlation structure between host and pathogen populations, producing false positive signals. Figure 1B shows that, in the absence of correction for stratification, 0.5% of association results (N=5,944) are false positive. However, both host or pathogen covariates are able to fully capture host or pathogen stratification. Therefore, using host covariate, pathogen covariate, or both, allows for a proper control of false positive signals, restoring a median p-value of 0.5. Details of this scenario are available in the online supplementary materials [30] by selecting ‘Optimal host and pathogen correction’.

**Fig. 1.**
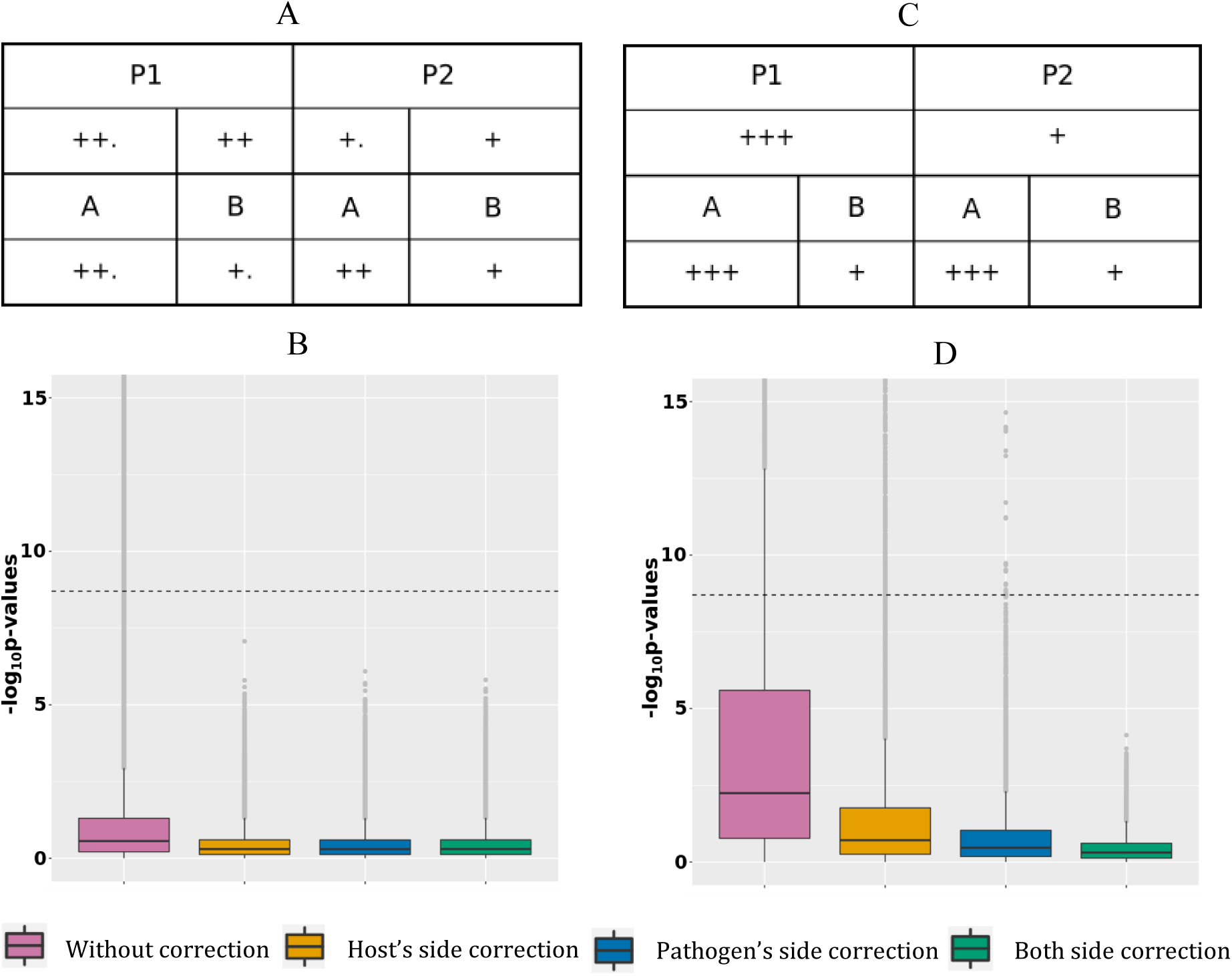
False positives signal. (A): Simulated host and pathogen variant frequency distribution for case 1. (B): P-value boxplot for case 1. (C): Simulated host and pathogen variant frequency distribution for case 2. (D): P-value boxplot for case 2.

**Scenario 2, suboptimal correction:** This scenario is represented in figure 1C, with 39,900 tests between 200 host SNPs (line 3 and 5 of Table 1) and 200 pathogen variants (line 2 and 4 of Table 2). 100 tests are on pairs in a causal relation and will be discussed in the next section. Host SNPs are stratified at two levels: major stratification between P1 and P2 and minor stratification between hosts infected by the two pathogen groups (resulting in higher allele frequencies for hosts infected by pathogen A, regardless of population, (figure 1D, additional ‘.’ for host populations infected by A). Pathogen variants are also stratified at two levels: major stratification between A and B and minor stratification between pathogens present in the two host populations (resulting in higher pathogen allele frequencies in P1, regardless of the pathogen group, figure 1D, additional ‘.’ for pathogen groups within P1).

False positive associations between host SNPs and pathogen variants can here result from: [A] major stratification on both sides (similar to scenario 1, above); [B] the correlation structure between major and minor stratification; and [C] the correlation structure between minor stratification on both sides. Figure 1D shows that not adjusting for stratification produces 9.8% (N=2,954) of false positive associations with a median p-value of 5.7 × 10-^3^. Because the minor stratification is too weak to be entirely captured by principal components, using host covariate cannot entirely correct for stratification and leaves 1.4% (N=409) of false positive associations with a median p-value of 1.9 × 10^−1^. Similarly, using only the pathogen covariate does not capture the minor stratification and leaves 0.07% (N=23) of false positive associations with a median p-value of 3.4 × 10-^1^. Finally, including both covariates allows for a proper control and restores a median p-value of 0.5. Therefore, correcting for both host and pathogen stratification prevents false positive associations. Details of this scenario are available in the online supplementary materials [30] by selecting ‘Suboptimal host and pathogen correction’.

##### 3.1.3 Power gain

This scenario presents association tests between 100 pairs of associated host SNPs and pathogen variants. The causal association is represented by the ‘arrow’ sign from host to pathogen (figure 2A).

**Fig. 2.**
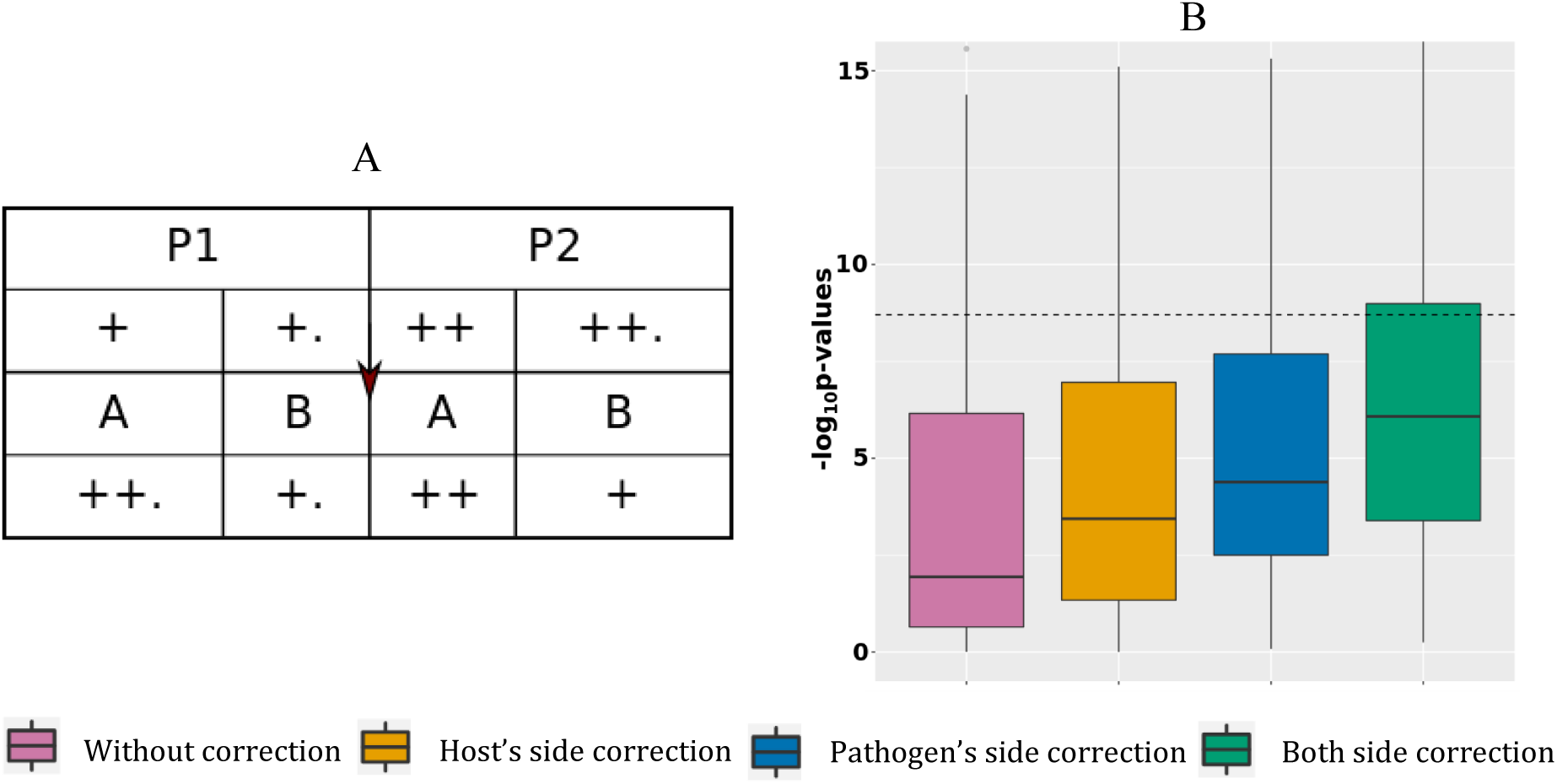
Power gain. (A): Simulated data structure for stratified host and pathogen data with true associations. (B): P-value boxplot for stratified host and pathogen data with true associations.

For this scenario, the host SNPs (l.5 Table 1) are stratified at two levels: major stratification between P1 and P2 (with higher allele frequencies in P2 than P1), and minor stratification between hosts infected by the two pathogen groups, resulting in higher allele frequencies for hosts infected by pathogen B.

To generate the pathogen variants (l.4 Table 2) for this scenario we used equation 3 and 4, described in section 2.2.2. This requires the host SNPs, binary background variations for the pathogen (*Y*_*bg*_) and another set of binary pathogen variations, associated with the SNPs (*Y*_*ca*_). The pathogen background variations or *Y*_*bg*_ are generated randomly, with stratification at two levels: major stratification between A and B (with higher pathogen allele frequencies in A than in B), and minor stratification between pathogens present in the two host populations, resulting in higher pathogen allele frequencies in P1. This distribution of pathogen background variation, along with the distribution of host variations, results into a negative correlation between host and pathogens because of two reasons. First, because the host population with a lower alternate allele frequency has pathogen genomes with a higher mutation rate, and vice versa. Second, because P1 and P2 infected by pathogen group B (lower mutation frequency) have a slightly higher alternate allele frequency than P1 and P2 infected by A (higher mutation frequency).

To generate the causal associations or *Y*_*ca*_, we used equation 3 in section 2.2.2. This makes *Y*_*ca*_ positively correlated with the host SNPs as the samples with fewer SNPs also have lower mutation rates in *Y*_*ca*_. Using these positively correlated *Y*_*ca*_ and negatively correlated *Y*_*bg*_ for generating the final pathogen variations (equation 4, section 2.2.2), nullifies the correlation structure due to stratification. This results in host and pathogen datasets in which causal associations are hidden under random noise due to stratification.

In figure 2B, without correction the median p-value is 0.11 with 16% of significant associations. Correcting the model using host covariates decreased the median p-value to 3.6 × 10-^4^ (20% of significant associations), while adjusting the model using pathogen covariate decreased the median p-value to 4.1 × 10-^5^ (21% of significant associations). Finally, adjusting the model using both host and pathogen covariates decreased the mean p-value to 8.3 × 10-^7^, with 29% of significant associations. These results show that correcting for stratification on both host and pathogen sides reduces the number of false negative signals. The mean p-value from the model corrected for both host and pathogen stratification is comparable to what we observed in the case with association and no stratification (mean p-value of 6.7 × 10-^7^), with an expected minimal power loss due to stratification of associated SNPs. Details of this scenario are available in the online supplementary materials [30] by selecting ‘Power gain’.

#### 3.2 Joint analysis of human and HIV genetic variation

To compare the results of our simulations to real-life data, we accessed human and viral genomic data collected from 1668 HIV-1 infected individuals participating in the Swiss HIV Cohort Study. We purposely selected a heterogeneous sample to ensure human and viral stratification. The principal component analyses showed diversity in terms of ethnicity, with 88% Caucasians, 5% Asians and 4% Africans (figure 3A); and of HIV-1 subtypes, with 81% of subtype B and 15% of subtype C infections (figure 3B).

**Fig. 3.**
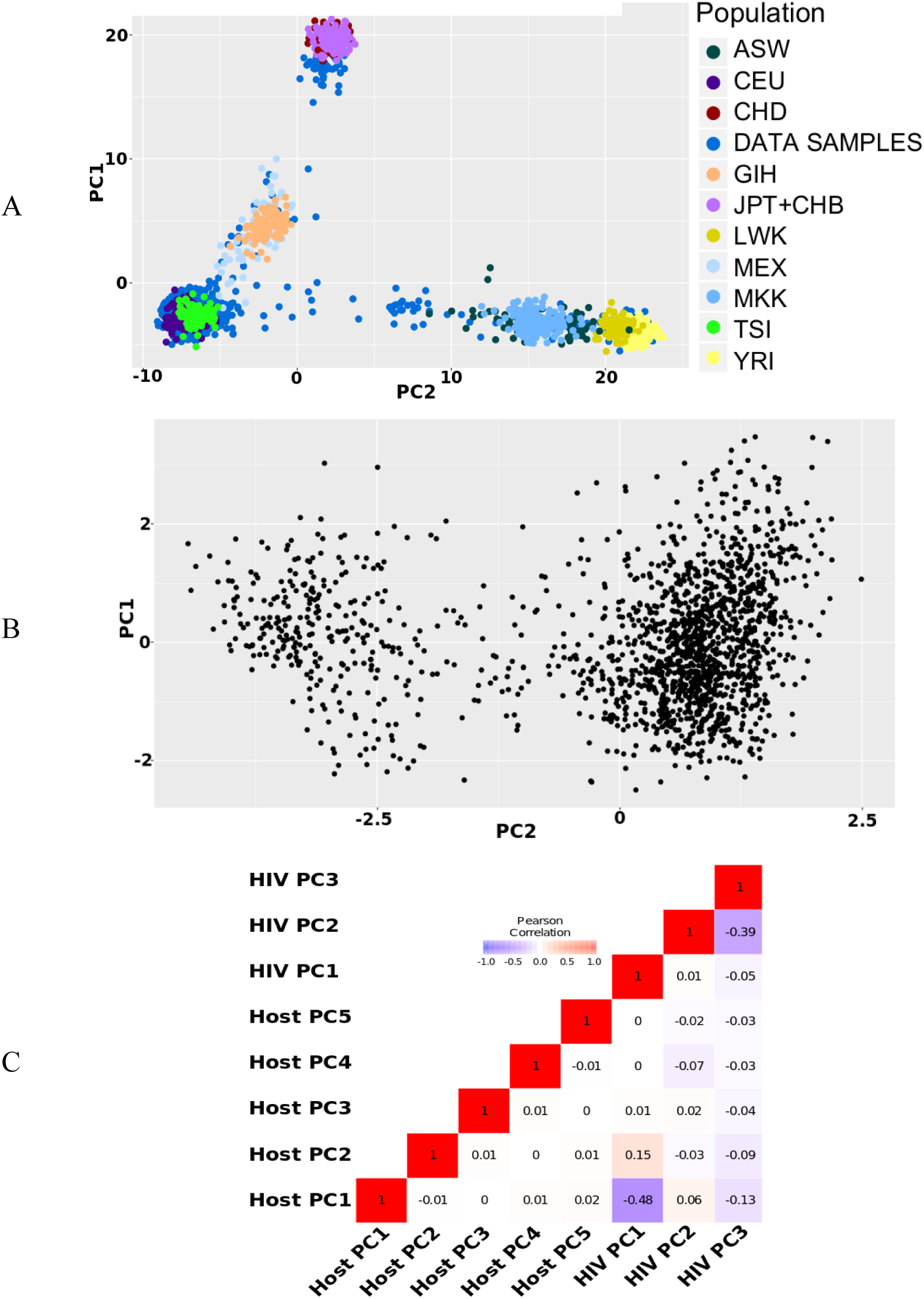
Population structures in HIV data. (A): Principal component plot for host data (first and second axis) (B): Phylogenetic principal component plot for the HIV virus data (first and second axis) (C): Pearson correlation between first five host principal components and first three HIV virus phylogenetic principal components

The human genome-wide data consisted of 5,600,166 SNPs, obtained after quality control, imputation and filtering. Binary HIV amino acid variants (N=403) were obtained from sequence data in the protease (N=155, 40 amino acid positions) and reverse transcriptase (N=248, 82 amino acid positions).

We searched for associations between individual SNPs and amino acid variants using logistic regression. To correct for population stratification on the human side, we included the top five principal components of the genotyping data in the regression models. The principal component plot in figure 4A shows the clustering of study samples with European, Asian and African reference samples from the HapMap dataset.

**Fig. 4.**
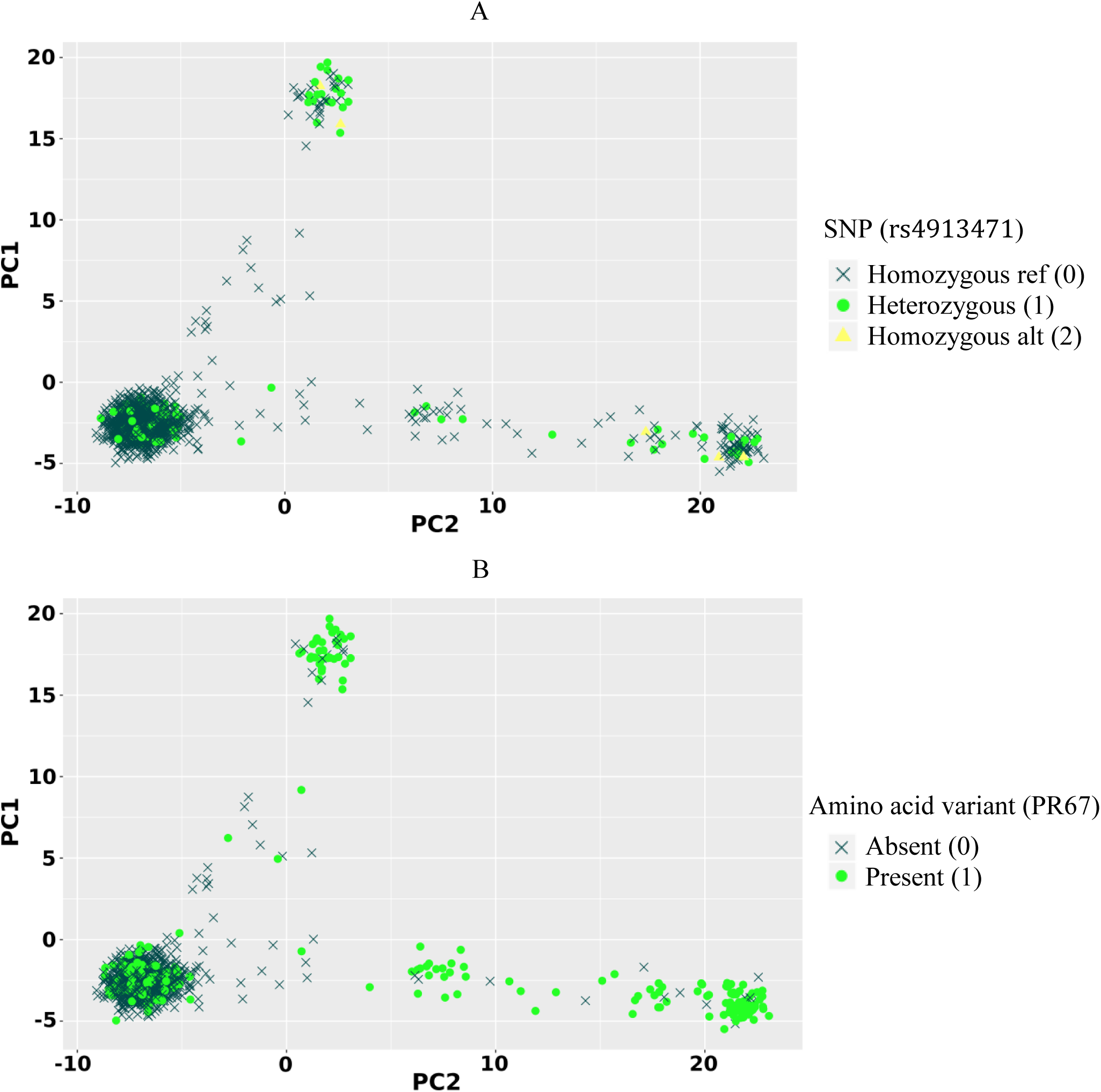
Allelic distribution of host SNP rs4913471 and HIV amino acid variant at position 67 in the protease region. (A): Genotypes of rs4913471 plotted on first two host principal components. (B): Presence or absence of amino acid variant at position 67 in the protease region plotted on first two host principal components.

To correct for pathogen stratification, we included the top three phylogenetic principal components (pPCs) [31] calculated using a phylogenetic tree built from the assembled HIV sequence data. Phylogenetic principal components control for phylogenetic covariance while producing PCA-like ordination. Figure 4B shows that the HIV genomes clustered mainly into two large groups (the first pPC explained 30% of the variation), on the first two pPC axes. We also computed Pearson correlation between the top five host principal components and the top three viral pPCs. Figure 4B shows only a weak correlation between host PCs and HIV pPCs, suggesting a need to correct for both to efficiently handle human and viral stratification.

Similar to what we did in the simulation study, we compared the association p-values obtained with four correction approaches. Upon correction for both host and pathogen stratification, we observed multiple highly significant associations between SNPs in the major histocompatibility complex (MHC) region of chromosome 6 and HIV amino acid variants. After correction for multiple testing (*p* < 2.2 × 10^−11^), significant associations were observed with 9 positions in the HIV proteome (5 in the PR region and 4 in the RT region). The strongest association was between rs2844527 and RT position 135 (*p* = 3.4 × 10-^35^). The top associated SNP for each HIV position is listed in Table 3. We replicated the associations, detected by Bartha et al. [13], for the protease position 35 (associated SNPs in complete LD with rs2523577), 93 (associated SNPs in complete LD with rs2263323) and the reverse transcriptase position 135 (associated SNPs in high LD with rs1050502 with = 0.7). In addition, we found new associations between SNPs in the MHC region and multiple other positions in the protease and the reverse transcriptase regions.

**Table 3.** HIV G2G associations

| HIV Gene | Position (amino acid residue) | SNP (Chromosome: bp) | P-value |
| --- | --- | --- | --- |
| PR | 35(E) | rs2596477 (6:31327723) | 8.503e-28 |
|  | 35(D) | rs17199328 (6:31322395) | 4.083e-25 |
|  | 12(T) | rs75344417 (6:31429439) | 1.559e-15 |
|  | 12(A) | rs116855165 (6:31059097) | 1.266e-11 |
|  | 37(N) | rs2596477 (6:31327723) | 3.370e-14 |
|  | 93(I) | rs9378249 (6:31327701) | 1.953e-13 |
|  | 93(L) | rs9378249 (6:31327701) | 1.953e-13 |
|  | 67(Y) | rs9391775 (6:31427948) | 3.394e-12 |
| RT | 135(I) | rs2844527 (6:31367636) | 3.444e-35 |
|  | 135(T) | rs79556279 (6:31329846) | 1.114e-29 |
|  | 165(T) | rs2442724 (6:31319907) | 1.557e-13 |
|  | 165(I) | rs92647589 (6:31248434) | 1.919e-12 |
|  | 123(E) | rs114773933 (6:31148349) | 1.917e-12 |
|  | 138(A) | rs114073761 (6:31336749) | 1.929e-11 |

As expected, correcting for population stratification at the human and/or viral levels led to an improvement in association p-values for known associated variants (“true positives”). For example, the p-value for the association between rs9266628 in the MHC region and HIV position 135 (presence or absence of amino acid residue I) in the reverse transcriptase region was 5.2 × 10-^18^ with no correction for stratification, 1.1 × 10-^18^ with correcting for host stratification only, 5.9 × 10-^19^ with correcting for pathogen stratification only, and 3.7 × 10-^20^ with correction for both host and pathogen stratification.

Conversely, several false positive associations (type I errors) could be identified. For example, the p-value for the association between rs4913471 on chromosome 12 and HIV position 67 (presence or absence of amino acid residue T) in the protease region was 1.5 × 10-^16^ with no correction for stratification, 3.2 × 10-^7^ with correcting for host stratification only, 4.5 × 10-^5^ with correcting for pathogen stratification only, and 3 × 10-^4^ with correcting for both host and pathogen stratification. To show the absence of true association, we plotted the distribution of genotypes for rs4913471 as well as the distribution of 1s and 0s for the HIV amino acid, over the host principal components (figure 4). Ethnicity correlates with both the SNP allele frequency (4A) and the presence or absence of the amino acid residue (4B), making this a classical case of false positive association due to stratification.

The summary statistics of this G2G analysis for all positions presented on Table 3 are available online [32].

## 4 Discussion and Conclusion

Genome-to-genome (G2G) association analyses can help dissect complex interactions between host and pathogen by integrating their respective genomic variation in a single model. The allele frequency distribution of genetic variants is not uniform in populations, resulting in systematic differences (“stratification”) upon sampling of host and pathogen populations. Such population stratification, if not accounted for properly, can lead to false positive association signals as well as false negative results. We here performed a comprehensive analysis to understand the various aspects of population stratification correction in a G2G framework.

Our simulation study demonstrated three main points that characterize the stratification correction effects. First, in the absence of stratification, the inclusion of covariates to correct for potential stratification only has a negligible impact on the results. Second, the existence of stratification on both sides produces false positive signals, which can be best minimized by including covariates that summarize both host and pathogen stratification. Third, population stratification can weaken true association signals, resulting in false negative results. In this case, correcting for stratification increases power and facilitates the identification of true associations.

We also presented a framework to properly correct for pathogen stratification by using pPCs, derived from the pathogen phylogenetic tree, as covariates. In our joint analysis of human and HIV genetic variation, we corrected for stratification using pPCs for HIV and principal components from the human genotyping data. We identified strong association signals between several SNPs in the MHC region and amino acid variants in the HIV proteome. Several of them were reported in a previous HIV G2G analysis, which was performed in a very homogeneous human population and only corrected for population stratification on the viral side.

In summary, we show that correcting for both host and pathogen stratification is necessary for unbiased G2G analysis. We also provide a framework that can adjust for stratification in the absence of reliable categorical labels, by exploiting host and pathogen genetic information. Our simulation design is implemented in R and available via GitHub.

## Members of the Swiss HIV Cohort Study

Anagnostopoulos A, Battegay M, Bernasconi E, BÖni J, Braun DL, Bucher HC, Calmy A, Cavassini M, Ciuffi A, Dollenmaier G, Egger M, Elzi L, Fehr J, Fellay J, Furrer H (Chairman of the Clinical and Laboratory Committee), Fux CA, Günthard HF (President of the SHCS), Haerry D (deputy of “Positive Council"), Hasse B, Hirsch HH, Hoffmann M, HÖsli I, Huber M, Kahlert C, Kaiser L, Keiser O, Klimkait T, Kouyos RD, Kovari H, Ledergerber B, Martinetti G, Martinez de Tejada B, Marzolini C, Metzner KJ, Müller N, Nicca D, Paioni P, Pantaleo G, Perreau M, Rauch A (Chairman of the Scientific Board), Rudin C (Chairman of the Mother & Child Substudy), Scherrer AU (Head of Data Centre), Schmid P, Speck R, StÖckle M, Tarr P, Trkola A, Vernazza P, Wandeler G, Weber R, Yerly S.

